# gpuZoo: Cost-effective estimation of gene regulatory networks using the Graphics Processing Unit

**DOI:** 10.1101/2021.07.13.452214

**Authors:** Marouen Ben Guebila, Daniel C Morgan, Kimberly Glass, Marieke L. Kuijjer, Dawn L. DeMeo, John Quackenbush

## Abstract

Gene regulatory network inference allows for the study of transcriptional control to identify the alteration of cellular processes in human diseases. Our group has developed several tools to model a variety of regulatory processes, including transcriptional (PANDA, SPIDER) and post-transcriptional (PUMA) gene regulation, and gene regulation in individual samples (LIONESS). These methods work by performing repeated operations on data matrices in order to integrate information across multiple lines of biological evidence. This limits their use for large-scale genomic studies due to the associated high computational burden. To address this limitation, we developed gpuZoo, which includes GPU-accelerated implementations of these algorithms. The runtime of the gpuZoo implementation in MATLAB and Python is up to 61 times faster and 28 times less expensive than the multi-core CPU implementation of the same methods. gpuZoo takes advantage of the modern multi-GPU device architecture to build a population of sample-specific gene regulatory networks with similar runtime and cost improvements by combining GPU acceleration with an efficient on-line derivation. Taken together, gpuZoo allows parallel and on-line gene regulatory network inference in large-scale genomic studies with cost-effective performance.

gpuZoo is available in MATLAB through the netZooM package https://github.com/netZoo/netZooM and in Python through the netZooPy package https://github.com/netZoo/netZooPy.

## INTRODUCTION

Gene regulation plays an important role in defining cell phenotypes and controlling cellular functions (1). Transcription factors (TFs) are regulatory proteins that bind promoter and enhancer regions near a gene to control its transcription and, ultimately, to mediate cellular processes (2,3). Several methods have been developed to infer gene regulatory networks from gene expression data and other data types (4-7). PANDA (Passing Attributes between Networks for Data Assimilation) (8,9) is an algorithm that estimates gene regulatory networks that are comprised of the collection of interactions between transcription factors and their target genes. Calculating such networks for individual phenotypes allows us to compare networks between phenotypes and understand changes in regulatory processes linked to health and disease. PUMA (10) estimates miRNA regulation (1) by seeding PANDA with miRNA estimated targets, while SPIDER (11) integrates DNase-seq data into the PANDA process to improve the accuracy of the network by restricting TF binding to open chromatin regions. LIONESS (12) makes iterative calls by embedding aggregate network reconstruction approaches such as PANDA or PUMA, in a loop and uses linear interpolation to calculate sample-specific gene regulatory networks for each member of a study population. Computing sample-specific networks informs us about gene regulation in various biological states by measuring heterogeneity within a population. These methods have proven quite useful, providing insight into, for example, tissue-specific gene regulation (13), explain sex-specific response to cancer drugs (14), and altered pathways in ovarian cancer (15).

Despite the success of these methods, one factor limiting the broader use of PANDA, PUMA, SPIDER, and LIONESS is the considerable memory space and computational time these algorithms require. As detailed below, PANDA, PUMA, SPIDER, and LIONESS perform a large number of matrix operations that were, until recently, reliant on CPUs composed of a relatively small number of computing cores that can maximally account for a few simultaneous software threads.

Graphics processing units (GPUs) offer an attractive alternative to CPUs, handling these repetitive matrix calculations in a faster and more efficient fashion. GPUs have hundreds of cores designed to handle many threads and thus support the efficient implementation of highly parallel computation in genomics (16) and in network inference (5). Since PUMA and SPIDER run on the same computational backend as PANDA, we will refer to the implementation of PANDA, PUMA, and SPIDER on the GPU as gpuPANDA. Therefore, gpuZoo consists of gpuPANDA which is a fast implementation of PANDA optimized to take advantage of the GPU architecture, and gpuLIONESS that implements LIONESS on multi-GPU devices to parallelize the required iterative computation of sample-specific networks. A cost-performance analysis found gpuPANDA to be up to 61 times faster and 28 times less expensive than running multi-threaded CPU implementations of PANDA, with similar performance improvements for gpuLIONESS of about 10x speedup for network modeling on a population of 127 individuals.

## MATERIAL AND METHODS

### The serial implementation of PANDA, PUMA, SPIDER, and LIONESS

Each cell contains proteins called transcription factors (TFs) that bind to specific DNA sequences to regulate gene expression. These transcription factors often work together to collectively regulate gene expression (3). These interactions can be represented in networks. We also know that genes that are regulated by the same transcription factors generally display similar patterns of expression. To capture these interactions, PANDA takes as input three “seed” matrices representing the networks of potentially “interacting” and co-regulated elements.

The first of these is a transcription factor-by-gene regulatory matrix (*W*_0_) that can be constructed by connecting TFs to their target genes based on mapping each TF’s known regulatory motif to the genome and identifying transcription factor binding sites (TFBS) within a window surrounding the transcription start site of each gene. This is based on our understanding that TFs often regulate gene expression by binding to the promoter region of their target genes.

The second matrix is a TF-by-TF cooperativity network (*P*_*0*_) that is based on “protein-protein interaction” (PPI) data collected from various sources such as *in vitro* experiments, text mining, and computational inference. Therefore, PPI data consists of pairs of proteins that interact with each other, for example, through physical binding. This matrix represents the network of proteins that may interact with each other to form multi-protein, i.e., multi-TF, complexes that together regulate specific genes. The use of PPI data in PANDA allows the algorithm to consider both direct regulation mediated by TFs binding to motifs on the DNA as well as indirect regulation by TFs that bind to other TFs that themselves bind to the DNA.

The final input matrix is the expression co-regulatory matrix (*C*_*0*_). The elements of this gene-by-gene matrix are pairwise Pearson Correlation Coefficients (PCC) between the corresponding gene pairs. PANDA integrates this correlation network with *W*_0_ based on the hypothesis that genes co-regulated by the same transcription factors should have correlated patterns of gene expression.

Because the regulatory network (*W*_0_), the cooperativity network (*P*_*0*_), and the co-regulatory matrix (*C*_*0*_) have different scales, the entries of each are Z-score standardized across both rows and columns. PANDA then iteratively optimizes the consistency between the three input matrices. It first calculates “Responsibility” and “Availability” values for each TF-gene edge and combines these values to update *W*. Next, it updates the values in *P* and *C*. Each of these updates uses a function based on a modified Tanimoto similarity for continuous variables, which we refer to as the Tfunction; the Tfunction can be conceptualized in terms of large matrix operations, making it amenable to significant improvement using GPU computing (see below and Supplementary methods).

PANDA (8) computes a final regulatory network (*W*_*f*_) (Figure S1-B) using a step-wise approach defined by a learning rate (α) (Supplementary methods). To better interpret performance gains from GPU computing, in addition to the Tfunction, we also included seven commonly used similarity metrics (Euclidean, squared Euclidean, standardized Euclidean, City block, Chebychev, Cosine, and PCC) as alternatives for benchmarking purposes (see Supplementary methods).

PANDA has recently been extended to incorporate additional regulatory mechanisms. PUMA (10) estimates the regulation of target genes by miRNAs by seeding a modified version of the PANDA algorithm with an estimate of miRNA target predictions in the *W*_0_ matrix. SPIDER (11) improves the accuracy of PANDA networks by integrating DNase-seq data to identify open chromatin regions where TFs are likely to bind. This is done by filtering *W*_0_ edges to the ones where the chromatin is in the open state.

In addition to aggregate methods that compute a context-specific network using several gene expression samples, we developed LIONESS (12) as an algorithm that uses linear interpolation to estimate single-sample networks. LIONESS first calculates a model (*W*) for the entire population of *N*_*S*_ samples. Then, it iteratively leaves out single samples, calculates a model for the population deprived of the *i*^th^ sample (*W*^*(i)*^), and uses the difference between these two models to estimate the network for the *i*^th^ sample (*W*^*i*^) using equation 4 in (12).

In our previous work, we have applied LIONESS to aggregate network models calculated using PANDA (14,17,18). In this case, computing a LIONESS network requires the following steps:

1. Compute a PANDA network (*W*) for all samples using PPI, motif, and gene coexpression. These three networks are normalized as a preprocessing step.
2. For a given sample *i*, compute gene coexpression using all samples but sample *i*, then normalize this matrix. PANDA is called on the newly obtained gene coexpression, motif, and PPI networks.
3. A GRN for sample *i* is derived by linear interpolation (12).
4. Step 2-3 are repeated for all the samples in the gene expression dataset.

In general, the slowest step in this process is computing and normalizing gene coexpression for every sample. However, since we are only interested in computing gene coexpression deprived of sample *i*, we can optimize the algorithm by computing gene coexpression on-line, i.e., by inferring the sample-deprived gene coexpression *C*^(*i*)^ from computing three initial variables: *m*, a vector representing the mean expression of genes across all samples; *s*, a vector representing the standard deviation in the expression of genes across all samples; and *Cov*, a matrix representing the covariance in expression between pairs of genes across all samples.

First, we use *m* to compute a vector representing the mean expression of genes across all samples except for sample *i*:

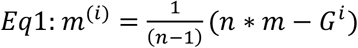

with *n* the number of samples and *G*^*i*^ the expression of genes in sample *i*. Next, we use *s* and *m*^*(i)*^to compute the standard deviation of genes across all samples except for sample *i*:

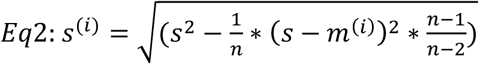

Third, we use *Cov* and *m* to compute the covariance matrix across all samples except for sample *i*

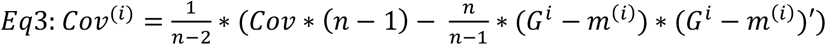

Finally, the sample-deprived co-expression matrix can be computed as follows:

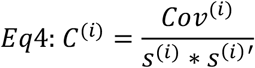

Computing the mean, standard deviation, and covariance only one time in step 1 allows us to infer the coexpression matrix for all samples in step 2. It avoids having to compute gene co-expression estimates independently for hundreds of samples.

### gpuPANDA and gpuLIONESS

Briefly, gpuPANDA implements network inference in parallel by first broadcasting data matrices to the GPU device and then performing all of the computation steps of the algorithm in the GPU device by distributing element-wise matrix operations, such as additions and multiplications (Figure S1-A), across hundreds of GPU cores using CUDA (19). For example, to determine the potential for a regulatory interaction between a TF and gene, PANDA computes the similarity *t(x,y)* between the target profile of the TF and the coexpressed partners of the target gene, as represented by in the matrices *W* and *C*. For each TF and gene pair, represented by row *x* of *W* and column *y* of *C*, the similarity is computed as follows:

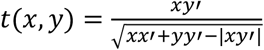

In CPU, the similarity between each TF-gene pair is computed sequentially. In GPU, each core computes the similarity for a single TF-gene pair. This means that the GPU/CPU speedup factor can be, in theory, of the order of the number of GPU cores (Figure S1-A), although in reality, GPU cores are much slower than CPU cores, and CPU cores can be multithreaded. In the end, the resulting network is sent back from the GPU to local memory and reduced to the final result (Figure S1-B). The gpuPANDA implementation has additional features such as the option to optimize GPU memory by considering only half of symmetrical coexpression matrices. In order to avoid memory transfer overhead, communication with the CPU was reduced to the case of device failure to save intermediary results and restart from the last iteration.

gpuLIONESS, which is essentially a series of batch calls to another network reconstruction approach, such as gpuPANDA, takes advantage of the architecture of multi-GPU devices, such as the NVIDIA TESLA K80 and NVIDIA TESLA P100, by assigning the computation of each single-sample network to an individual GPU device in parallel. gpuLIONESS uses the MATLAB and Python interfaces to embed CUDA (19) processes for each NVIDIA GPU in a Message Passing Interface (MPI) process (20) to compute single-sample networks in parallel. This hybrid structure provides two levels of parallelism that ensures message passing of computation results between non-shared memory processes and within each CUDA process.

gpuZoo which consists of gpuPANDA and gpuLIONESS was implemented in MATLAB (2019a, version 9.6.0, The MathWorks Inc., Natick, Massachusetts, USA) as part of the netZooM package (https://github.com/netZoo/netZooM; version 0.5.2) and in Python (version 3.7) as part of the netZooPy package (https://github.com/netZoo/netZooPy; version 0.6.2).

### Benchmarking procedure

The runtime and cost of network generation for the CPU and GPU implementations of PANDA and LIONESS were compared using networks of three sizes: 652 TFs by 1000 genes, 652 TFs by 27,149 genes, and 1603 TFs by 43,698 transcripts. These roughly correspond to the sizes of a small network, protein-coding genes network, and transcript network, respectively.

The small size network was derived from the input data used by Lopes-Ramos and colleagues (21) to construct lymphoblast cell line (LCL) regulatory networks using i) expression data from GTEx (22), ii) PPI data from STRINGdb (23), and iii) TF binding predictions derived using FIMO (24) to scan the promoter regions of all gene sequences defined as TSS +/-750bp in the human genome (hg38) for matches to human PWMs from CIS-BP (3). To create the small network from these data, we restricted the TF binding network to the first 1000 genes. In the data pre-processing step, we took the intersection of these three input data sources, i.e., the intersection of the TFs in PPI and TF binding motif matrices and the intersection of the genes in the gene coexpression and the TF binding motif matrices; this resulted in *W, P*, and *C* matrices that included data for 652 TFs and 1000 genes.

The protein-coding gene network was also derived from GTEx LCL cell line data, but in the data pre-processing step we did not restrict to 1000 genes and used the union of the three complete input data sets, which still had 652 TFs but which increased the number of target genes to 27,149.

Finally, to test the maximal capacity of the GPU hardware, we computed a large network consisting of all the known TFs and individual gene transcripts. These individual transcripts reflect the alternative splicing process in which each gene can code for several transcripts. This model was called the transcript network. The transcript network was based on THP-1 Leukemic monocyte cell line (25), gene expression data from GEO (26) processed in ARCHS4 (27) to obtain transcript levels, a PPI network of 1603 TFs encoded in the human genome from STRINGdb (23), and the same set of TF binding predictions used in the protein-coding gene network. In the data pre-processing step, we used the union of the three data sources which resulted in a data set consisting of 1603 TFs and 43,698 transcripts.

In addition to the default Tfunction similarity metric and default learning rate (α=0.1), we ran PANDA using seven commonly used similarity metrics that can be computed on the GPU (Supplementary Material) and two additional learning rates (α=0.2 and α=0.3) and compared them in terms of computational speed and cost. Two reasons motivated the expansion to additional parameters. First, we wanted to show that GPU versus CPU results are consistent across different parameters. Second, although we successfully used the similarity metric Tfunction with a learning rate of 0.1 in earlier studies (13,14,21), the cost-effective acceleration provided by gpuPANDA enables the exploration of additional parameter combinations. Therefore, we wanted to make sure that performance gains were guaranteed beyond the standard parameter values.

We wanted to assess the runtime and cost performance of PANDA and gpuPANDA as well as LIONESS and gpuLIONESS, first using MATLAB and then Python implementations. We used two CPU configurations (compute optimized CPU1 and memory optimized CPU2) and three GPU configurations (NVIDIA TESLA V100-GPU1, NVIDIA P100-GPU2 and a smaller NVIDIA K80-GPU3); the configurations are shown in detail in Table 1. To reduce the number of comparisons, our approach was to benchmark the transcript model in GPU1 because it could not be loaded in other devices, and to benchmark the coding-genes model in GPU1 and GPU2 for the same reasons. Finally, we benchmarked the small model on GPU2 and GPU3 because with larger devices, the initialization time could exceed the computation time.

**Table 1.** Specification of the hardware units used for benchmarking. All the CPU cores are used to perform computations because MATLAB enables hyperthreading by default. EC2 cost corresponds to AWS On-Demand price. *p3dn.24xlarge has 8 Tesla P100 Tensor Core, the original cost of 31.218 was divided by 8 to estimate the cost of one unit.

| Unit | AWS reference | Price (\$/hr) | Manufacturer reference | Number of cores | Clock speed (Ghz) | Memory (GB) | Specific ations |
| --- | --- | --- | --- | --- | --- | --- | --- |
| CPU1 | c5d.12xlarge | 2.304 | 2nd generation Intel Xeon Scalable Processors | 48 | 3.6-3.9 | 96 | compute - optimize d |
| CPU2 | r5a.8xlarge | 1.808 | AMD EPYC 7000 | 32 | 2.5 | 256 | memory-optimize d |
| GPU1 | p3dn.24xlarge | 3.902 * | Nvidia Tesla V100 Tensor Core | 5120 | 1.53 | 32 | Largest GPU |
| GPU2 | p3.2xlarge | 3.06 | Nvidia Tesla P100 | 3584 | 1.19 | 16 | Large GPU |
| GPU3 | p2.xlarge | 0.9 | Nvidia Tesla K80 | 2496 | 0.52 | 12 | Smaller GPU |

All analyses were performed on Amazon Web Services (AWS) running the MATLAB (version 2019a) distribution in Ubuntu 18.04 and Windows 10 that enables MATLAB memory benchmarking, and Python (version 3.7). The cost was computed as the cost of the instance on AWS multiplied by the runtime in seconds, since AWS EC2 bills by second.

## RESULTS

We first ran the MATLAB implementations of PANDA and gpuPANDA on our three test networks using three learning rate values (α=0.1, α=0.2, α=0.3) with calculations in single and double precision; we also ran these methods with each of the eight similarity metrics.

For the small network that includes 652 TFs and 1000 genes, both PANDA on CPU1 and CPU2 and gpuPANDA running on the GPU2 and GPU3 platforms were able to infer gene regulatory network models that were identical to one another as determined by the absolute value of the largest difference (Figure S2). This was true using all eight similarity metrics and running in both single and double precision. However, gpuPANDA demonstrated significant advantages in both runtime (up to 7-fold; Table S1-2) and cost (up to 15-fold; Figure 1-A, Table S3-4). In comparison to PANDA on CPU1, the rate of decrease of gpuPANDA runtime outpaced the decrease in cost for both GPU2 and GPU3 (Figure 1-B). However, since small networks do not require large device memory, gpuPANDA was more efficient with the smaller GPU3 and provided a decrease in cost at a larger rate than the decrease in runtime in comparison to CPU2.

**Figure 1.**
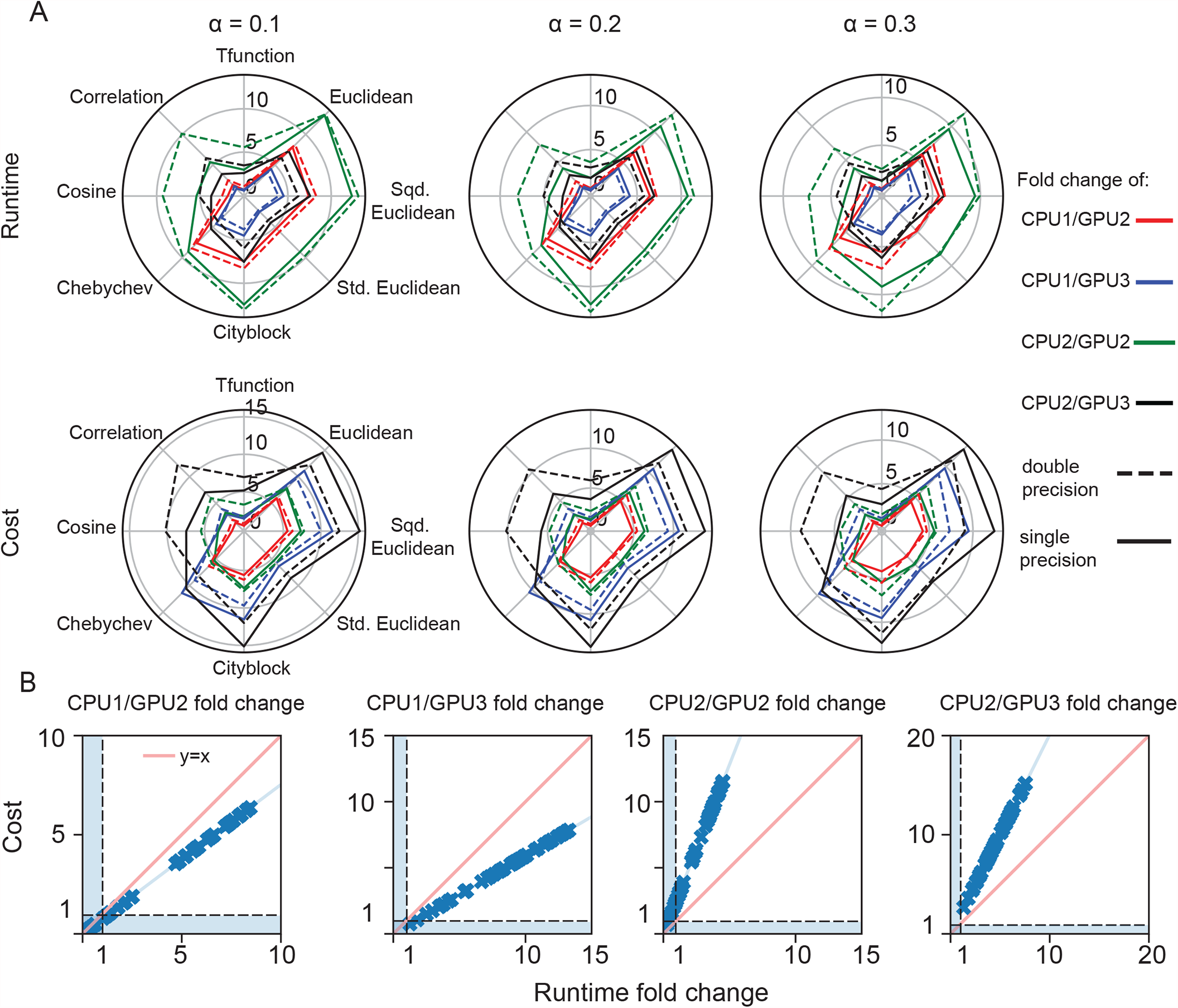
Runtime and cost performance of gpuPANDA in the small network. A- Runtime (first row) and cost (second row) fold change between CPU1, CPU2, GPU2, and GPU3. B-Rate of cost fold change as an effect of runtime fold change in small models in single and double precision. The blue area represents an increase in cost and/or runtime of GPU computation over CPU.

For the network modeled on protein-coding genes, GPU acceleration was possible in GPU1 but only in single precision with GPU2 due to memory limitations. gpuPANDA had up to ninefold decrease in runtime and sevenfold decrease in cost when comparing GPU2 and the compute-optimized device CPU1 (Figure 2-A). For the memory-optimized CPU2, the increase reached up to 26-fold for the runtime and we saw a decrease in cost of up to 15-fold (Figure 2-A). This was particularly clear with the modified Tanimoto (Tfunction) similarity metric at the default learning rate of 0.1. An analysis of cost fold change rates as a function of runtime fold change rates (Figure 2-B) showed that GPU2/CPU1 and GPU2/CPU2 performance growth evolved in a regimen where runtime decrease had a faster rate than the cost decrease. Similarly, GPU1 was up to 12 times faster than CPU1 and up 61 times faster than CPU2 particularly in double precision computation using the Euclidean distance, which corresponded to a 7-fold and 28-fold reduction in costs.

**Figure 2.**
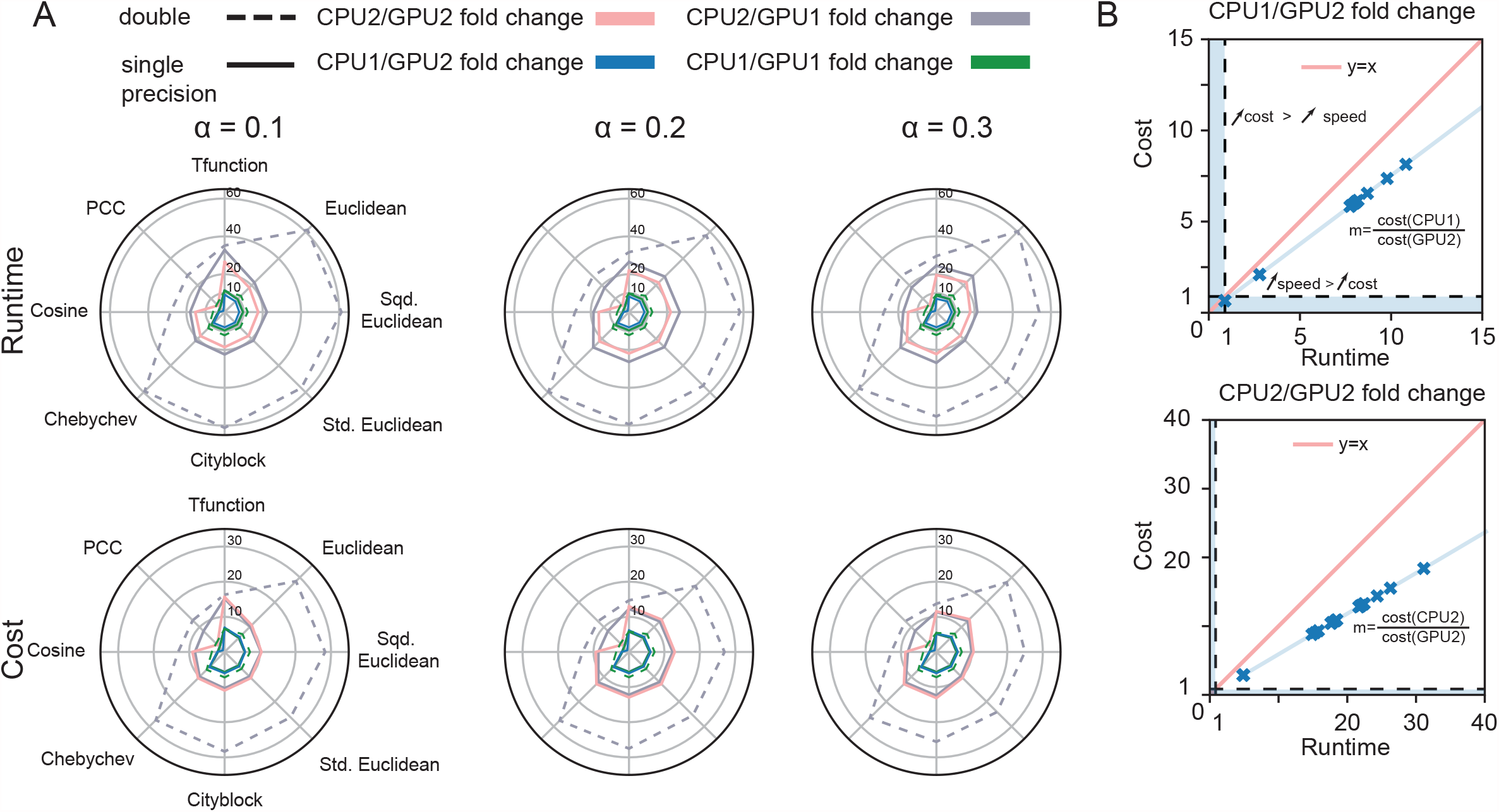
Runtime and cost performance of GPUs and CPUs on the protein coding-genes model. A-Fold change of runtime as a function of cost between CPU1 and GPU2 and CPU2 and GPU2 in single precision and for three values of learning rate (α). B- Effect of runtime fold change on cost fold change between CPU1 and GPU2 (top panel) and CPU2 and GPU2 (bottom panel).

We designed the GPU code to optimize memory usage. Specifically, we measured the memory requirements of PANDA and gpuPANDA across six sampling points after the function call (Figure 3-A) and found a 2.6-fold decrease in memory usage with the GPU implementation. However, despite this improvement, neither the GPU2 and GPU3 configuration had sufficient memory to load the input matrices of the transcript network and perform operations using either single or double precision (Figure 3-B). In addition, we could only load the network on GPU1 on single precision only for Tfunction similarity metric. Benchmarking against CPU1 and CPU2 in this setting revealed a 24-fold decrease in runtime (Figure 3-C) and 11-fold decrease in cost (Figure 3-D).

**Figure 3.**
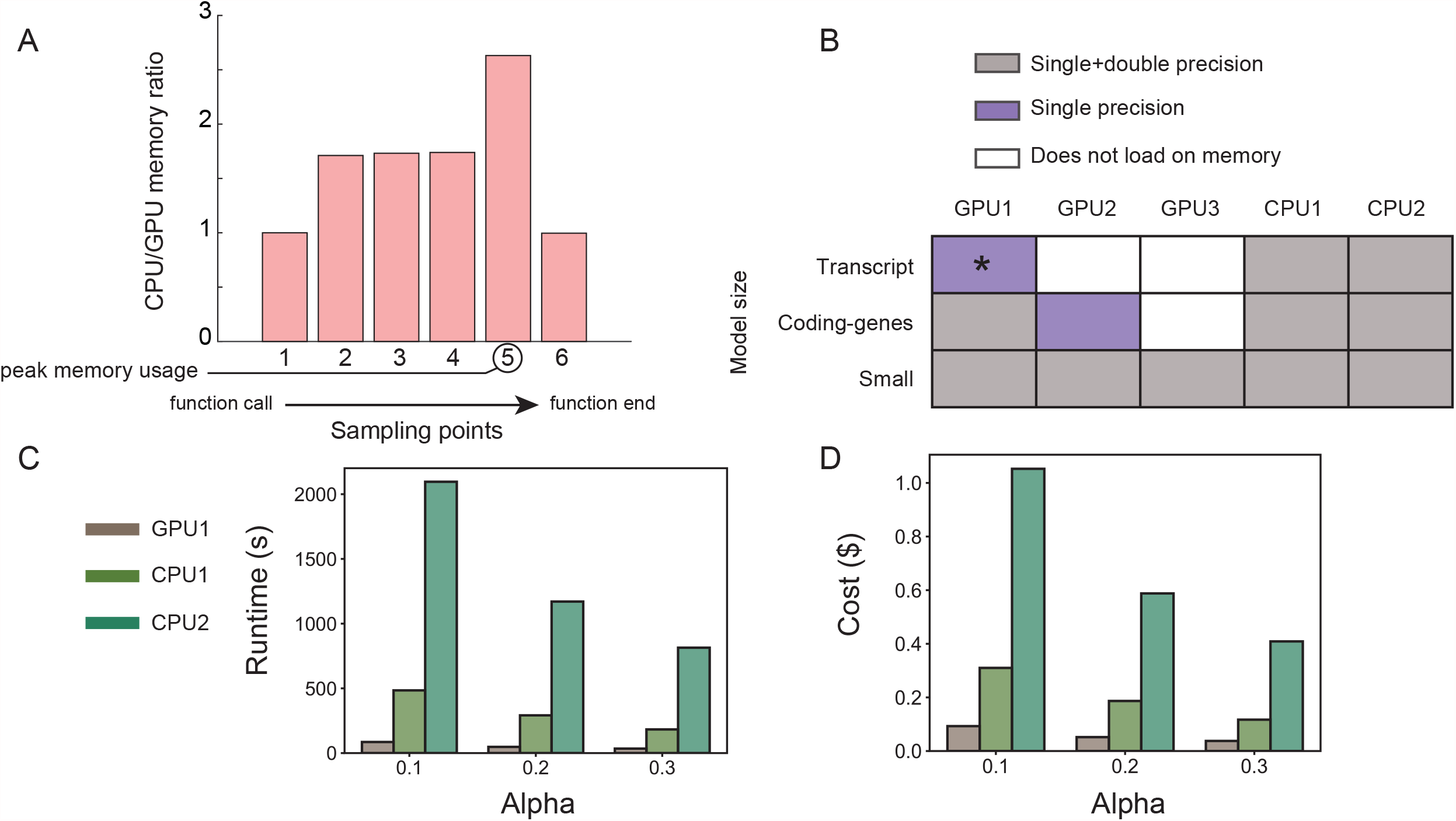
GPU performance on transcript model and memory benchmark. A-Memory usage of GPU implementation in comparison to CPU implementation. B-Precision capabilities for the tested hardware using the small network, protein-coding genes network, and transcript regulatory gene network. C-Runtime and D-cost of running transcript model on GPU1, CPU1, and CPU2 in single precision. *Single precision computation on GPU1 converges with Tfunction only.

In addition to testing the MATLAB code, we tested the Python implementations of PANDA and gpuPANDA on the small network. We found similar results to those described above that were based on the MATLAB implementation. For example, calculating single precision networks using PANDA on CPU2 and gpuPANDA on GPU3, gpuPANDA was more than 10 times faster than the CPU implementation. We also found the output networks to be identical, with the largest absolute difference in edge weights equal to 3.5×10^−5^ (Figure S3).

Finally, we tested LIONESS and gpuLIONESS in MATLAB. Both LIONESS and gpuLIONESS perform a series of batch calls, in our case to PANDA and gpuPANDA, respectively. In estimating 127 single precision individual sample networks based on the small network dataset, we found gpuLIONESS to be 10 times faster when running on GPU2 compared to LIONESS running on CPU2 and 24 times faster when running on GPU1 in the same configuration (Figure S4); the largest absolute difference between the CPU and GPU network edge weights was 0.015, which was less than 0.01% of that edge weight, while the average absolute difference was 6.5×10^−5^. We have also combined GPU speedup with an additional algorithmic improvement consisting of deriving the coexpression matrix on-line (Eq1-Eq4) without having to recompute it for every sample (Figure S4), however, this approach did not further increase the speedup.

## DISCUSSION

As the sample sizes for genomic and multi-omic data studies grow, we have the opportunity to develop increasingly accurate models of the potential causes of various diseases and phenotypic traits. However, the computational complexity, time, and cost of building such models has become a limiting factor in many applications. The development of PANDA, PUMA, SPIDER, and LIONESS as techniques for inferring accurate regulatory models has allowed the exploration of gene regulation in health and disease, but the use of these models has been limited by the availability of computational resources. For example, using PANDA and LIONESS to generate more than 9,435 individual sample networks (17) using data from GTEx v6 initially took multiple months running on a conventional multi-CPU cluster; rerunning those networks in response to a question from referees took more than six weeks (after having optimized the CPU code). Our interest in repeating this analysis with GTEx v8, and with other large datasets, underscores the need for additional computational improvements.

Fortunately, PANDA, PUMA, and SPIDER are implemented as a series of matrix operations that make them particularly amenable to implementation on GPUs (9). gpuPANDA represents an adaptation of these methods that parallelizes the large matrix operations in each iteration of the network inference and refinement process. The implementation of gpuLIONESS extends this further by distributing the calculation of PANDA networks for each leave-one-out data subset across the available GPU devices, such that the computation of each individual sample network is distributed across the cores available within each device.

Improving runtime was a major motivation for creating GPU implementations of PANDA and LIONESS. By taking advantage of Python and MATLAB GPU interfaces to CUDA (19), gpuPANDA reduced memory use by 2.6-fold relative to the CPU implementation, in part because it is able to take advantage of symmetries in the coexpression matrix (which is generally the largest matrix used in PANDA). We recognize that we might be able to further reduce memory usage by sending intermediate results to RAM to free space for the next iteration. However, we chose not to do so because the associated I/O would increase computation time and, consequently, cost.

Despite these improvements, neither of the GPU devices, including the larger memory GPU1 (32 GB), was able to load the data for the largest transcript network in double precision (43,698 transcripts and 1,603 TFs, Figure 3-D). This is not surprising, given the size of the coexpression matrix. However, it should not pose a major barrier to the use of gpuZoo since most network inference modeling only includes the 20,000-30,000 protein-coding genes. Additionally, most pipelines would further eliminate genes not expressed in a particular tissue during data preprocessing. In addition, the majority of our earlier investigations (13,14,17) fall within the size of the protein-coding genes network, for which the computations carried with the modified Tanimoto similarity (Tfunction) had the largest speedup with gpuPANDA. With GPU3, gpuZoo was not able to load the protein-coding genes network (652 TFs and 27,149 genes), and with GPU2 it was only able to load it in single precision. However, the loss of double precision in the matrix calculations does not produce major changes in the overall network estimation and likely has a much less significant effect than noise in measurements of gene expression (Figure S5).

Computing 127 single-precision, sample-specific networks using gpuLIONESS for the protein-coding genes network was 10-fold faster using GPU2. When inferring a sample-specific network using PANDA together with LIONESS, there is an additional step that requires recomputing and normalizing the gene co-expression matrix for each sample, which requires large memory resources due to the size of the matrix. When benchmarking GPU2 and due to memory constraints, this step was performed in CPU and then sent to GPU. In GPU1, this step could be performed on device, which improved the speedup to 24-fold. We also investigated combining GPU acceleration with computing gene coexpression on-line. We did not see an improvement in the total runtime in the context of our tested networks. However, we have investigated whether this approach could be beneficial when the number of samples increases relative to the number of genes. Running a comparison between coexpression and on-line coexpression on a 1,000 variable random matrix showed similar performance (Figure S6) when the number of samples was 0.5% the number of genes, which is about the ratio used in our study (127 samples and 27,149 genes). However, increasing the percentage indicated a 2.45 speedup when the number of samples was equal to number of genes, with benefits starting at 50% (1.5 speedup) and possibly even for lower ratios. In a recent work, we computed sample-specific gene regulatory networks using data from the Connectivity Map across 170,013 experiments on 12,328 genes in four days by combining acceleration from GPU and on-line coexpression. This would have required several weeks using classical approaches.

The largest absolute difference between the edge weights of single-sample gpuLIONESS networks was 0.015 which is larger than the difference between PANDA and gpuPANDA networks in single precision (∼1×10^−5^). However, the average error was equal to 6.5×10^−5^, which is within the order of single precision computation.

For the small network, despite a greater increase in network inference speed with GPU2, the smaller GPU3 was more cost-effective for a similar performance (Figure 2-A, Table S1, Table S3). In particular, computing gene regulatory networks using the similarity metric Tfunction on GPU2 was less efficient than GPU3 and CPU1, because initializing a large device requires more time than the computation itself.

Comparing inference of regulatory networks using gpuPANDA on three GPU architectures and PANDA on two CPUs, each with different specifications, allowed us to understand the effects of clock speed and memory access on the final runtime and cost. In particular, the CPU machines on which we ran PANDA were significantly different: CPU1, the compute-optimized machine has a faster processing speed but relatively limited RAM (96 GB) while the memory-optimized CPU2 has slower CPUs but far greater and faster accessible memory (256 GB) (Table 1).

The main drawback of these implementations is that they are unable to process networks with more than 20,000 genes in double precision. However, we found that the differences between single precision and double precision networks remain within the order of single precision, which indicates that neither hardware specifications nor the software implementation account for additional deviation in precision than what is expected (Figure S5). Therefore, computing in single precision when GPU memory is lacking could be a viable approach for networks that cover more than protein-coding genes.

Taken together, gpuZoo offers a fast and less computationally expensive option for the estimation of batches of gene regulatory networks. These implementations allow the estimation of gene regulation in large-scale genomics studies such as TCGA (28), the Connectivity Map (29), and the GTEx project (22). The fast development of GPU devices (30) will soon enable large-scale network inference in double precision. Finally, gpuZoo tools are enabling biological discovery by providing a computational engine that supports our recent endeavor to reconstruct gene regulatory networks across human conditions (31) (https://grand.networkmedicine.org).

## DATA AVAILABILITY

gpuZoo (gpuPANDA, gpuPUMA, gpuSPIDER, and gpuLIONESS) is available through the Network Zoo package (netZoo; netzoo.github.io) in MATLAB (netZooM v0.5.2) at https://github.com/netZoo/netZooM with a step-by-step tutorial https://github.com/netZoo/netZooM/tree/master/tutorials, and in Python (netZooPy v0.6.2) https://github.com/netZoo/netZooPy with a tutorial https://github.com/netZoo/netZooPy/tree/master/tutorials.

The code of the benchmarks is available at https://github.com/QuackenbushLab/gpuzoo, and the corresponding data is available at https://netzoo.github.io/zooanimals/gpuzoo/.

## SUPPLEMENTARY DATA

Supplementary data consists of one supplementary file and four supplementary tables.

## REFERENCES

1. Hobert, O. (2008) Gene regulation by transcription factors and microRNAs. Science, 319, 1785–1786.

2. Zeitlinger, J. (2020) Seven myths of how transcription factors read the cis-regulatory code. Current Opinion in Systems Biology.

3. Lambert, S.A., Jolma, A., Campitelli, L.F., Das, P.K., Yin, Y., Albu, M., Chen, X., Taipale, J., Hughes, T.R. and Weirauch, M.T. (2018) The Human Transcription Factors. Cell, 175, 598–599.

4. Irrthum, A., Wehenkel, L. and Geurts, P. (2010) Inferring regulatory networks from expression data using tree-based methods. PloS one, 5, e12776.

5. He, J., Zhou, Z., Reed, M. and Califano, A. (2017) Accelerated parallel algorithm for gene network reverse engineering. BMC systems biology, 11, 85–97.

6. Haury, A.-C., Mordelet, F., Vera-Licona, P. and Vert, J.-P. (2012) TIGRESS: trustful inference of gene regulation using stability selection. BMC systems biology, 6, 1–17.

7. Ruyssinck, J., Geurts, P., Dhaene, T., Demeester, P. and Saeys, Y. (2014) NIMEFI: gene regulatory network inference using multiple ensemble feature importance algorithms. PLoS One, 9, e92709.

8. Glass, K., Huttenhower, C., Quackenbush, J. and Yuan, G.C. (2013) Passing messages between biological networks to refine predicted interactions. PLoS One, 8, e64832.

9. Glass, K., Quackenbush, J. and Kepner, J. (2015), 2015 IEEE High Performance Extreme Computing Conference (HPEC). IEEE, pp. 1–6.

10. Kuijjer, M.L., Fagny, M., Marin, A., Quackenbush, J. and Glass, K. (2020) PUMA: PANDA Using MicroRNA Associations. Bioinformatics, 36, 4765–4773.

11. Sonawane, A.R., DeMeo, D.L., Quackenbush, J. and Glass, K. (2020) Constructing Gene Regulatory Networks using Epigenetic Data. bioRxiv.

12. Kuijjer, M.L., Tung, M.G., Yuan, G., Quackenbush, J. and Glass, K. (2019) Estimating Sample-Specific Regulatory Networks. iScience, 14, 226–240.

13. Sonawane, A.R., Platig, J., Fagny, M., Chen, C.Y., Paulson, J.N., Lopes-Ramos, C.M., DeMeo, D.L., Quackenbush, J., Glass, K. and Kuijjer, M.L. (2017) Understanding Tissue-Specific Gene Regulation. Cell Rep, 21, 1077–1088.

14. Lopes-Ramos, C.M., Kuijjer, M.L., Ogino, S., Fuchs, C.S., DeMeo, D.L., Glass, K. and Quackenbush, J. (2018) Gene Regulatory Network Analysis Identifies Sex-Linked Differences in Colon Cancer Drug Metabolism. Cancer Res, 78, 5538–5547.

15. Glass, K., Quackenbush, J., Spentzos, D., Haibe-Kains, B. and Yuan, G.C. (2015) A network model for angiogenesis in ovarian cancer. BMC Bioinformatics, 16, 115.

16. Taylor-Weiner, A., Aguet, F., Haradhvala, N.J., Gosai, S., Anand, S., Kim, J., Ardlie, K., Van Allen, E.M. and Getz, G. (2019) Scaling computational genomics to millions of individuals with GPUs. Genome biology, 20, 1–5.

17. Lopes-Ramos, C.M., Chen, C.Y., Kuijjer, M.L., Paulson, J.N., Sonawane, A.R., Fagny, M., Platig, J., Glass, K., Quackenbush, J. and DeMeo, D.L. (2020) Sex Differences in Gene Expression and Regulatory Networks across 29 Human Tissues. Cell Rep, 31, 107795.

18. Lopes-Ramos, C.M., Belova, T., Brunner, T., Quackenbush, J. and Kuijjer, M.L. (2021) Regulation of PD1 signaling is associated with prognosis in glioblastoma multiforme. bioRxiv.

19. Nickolls, J., Buck, I., Garland, M. and Skadron, K. (2008) Scalable Parallel Programming with CUDA. Queue, 6, 40–53.

20. Forum, M.P. (1994). University of Tennessee.

21. Lopes-Ramos, C.M., Paulson, J.N., Chen, C.Y., Kuijjer, M.L., Fagny, M., Platig, J., Sonawane, A.R., DeMeo, D.L., Quackenbush, J. and Glass, K. (2017) Regulatory network changes between cell lines and their tissues of origin. BMC Genomics, 18, 723.

22. Consortium, G.T., Laboratory, D.A., Coordinating Center -Analysis Working, G., Statistical Methods groups-Analysis Working, G., Enhancing, G.g., Fund, N.I.H.C., Nih/Nci, Nih/Nhgri, Nih/Nimh, Nih/Nida et al. (2017) Genetic effects on gene expression across human tissues. Nature, 550, 204–213.

23. Szklarczyk, D., Gable, A.L., Lyon, D., Junge, A., Wyder, S., Huerta-Cepas, J., Simonovic, M., Doncheva, N.T., Morris, J.H., Bork, P. et al. (2019) STRING v11: protein-protein association networks with increased coverage, supporting functional discovery in genome-wide experimental datasets. Nucleic Acids Res, 47, D607–D613.

24. Grant, C.E., Bailey, T.L. and Noble, W.S. (2011) FIMO: scanning for occurrences of a given motif. Bioinformatics, 27, 1017–1018.

25. Bosshart, H. and Heinzelmann, M. (2016) THP-1 cells as a model for human monocytes. Annals of translational medicine, 4.

26. Edgar, R., Domrachev, M. and Lash, A.E. (2002) Gene Expression Omnibus: NCBI gene expression and hybridization array data repository. Nucleic acids research, 30, 207–210.

27. Lachmann, A., Torre, D., Keenan, A.B., Jagodnik, K.M., Lee, H.J., Wang, L., Silverstein, M.C. and Ma’ayan, A. (2018) Massive mining of publicly available RNA-seq data from human and mouse. Nature communications, 9, 1–10.

28. Weinstein, J.N., Collisson, E.A., Mills, G.B., Shaw, K.R.M., Ozenberger, B.A., Ellrott, K., Shmulevich, I., Sander, C. and Stuart, J.M. (2013) The cancer genome atlas pan-cancer analysis project. Nature genetics, 45, 1113–1120.

29. Subramanian, A., Narayan, R., Corsello, S.M., Peck, D.D., Natoli, T.E., Lu, X., Gould, J., Davis, J.F., Tubelli, A.A., Asiedu, J.K. et al. (2017) A Next Generation Connectivity Map: L1000 Platform and the First 1,000,000 Profiles. Cell, 171, 1437–1452 e1417.

30. Bridges, R.A., Imam, N. and Mintz, T.M. (2016) Understanding GPU Power: A Survey of Profiling, Modeling, and Simulation Methods. ACM Comput. Surv., 49, Article 41.

31. Guebila, M.B., Lopes-Ramos, C.M., Weighill, D., Sonawane, A., Burkholz, R., Shamsaei, B., Platig, J., Glass, K., Kuijjer, M.L. and Quackenbush, J. (2021) GRAND: A database of gene regulatory network models across human conditions. bioRxiv.

