## Supplementary file for "gpuZoo: Cost-effective estimation of gene regulatory networks using the Graphics Processing Unit"

### I. Supplementary methods

#### 1. The PANDA algorithm

PANDA computes a regulatory network ($W_f$) using three inputs: An initial estimate of the regulatory network ($W_0$), a cooperativity network ($P_0$), and a co-regulatory matrix ($C_0$). The algorithm iteratively computes concordance between pairs of input matrices using a similarity metric (Figure S1-B). In the remainder of this section, we will refer to all similarity metrics by $S$.

First, the Responsibility is a function associated with the transcription factor. At each iteration *t*, it estimates the potential of a TF to regulate target genes in the network. We know that transcription factors often work cooperatively, forming complexes to regulate their target genes. Because PPI data should capture the potential interactions between TFs, PANDA uses a similarity metric to assay the degree of agreement between the TFs that target gene *j* ($W_{.j}^{(t)}$), and those that form a complex with TF *i* ($P_{i.}^{(t)}$). The Responsibility (*R*) is calculated as:

$$R_{ij}^{(t)} = S\left(P_{i.}^{(t)}, W_{.j}^{(t)}\right).$$

Second, we use a similar approach to calculate the Availability associated with each gene sharing an edge in $W_0$ with a TF, assuming that genes sharing similar patterns of expression across samples in a dataset are likely to be regulated by one or more of the same TFs. We test for agreement between the regulatory targets of TF *i* ($W_{i.}^{(t)}$) and the set of genes with which gene *j* has correlated expression ($C_{.j}^{(t)}$) to calculate the Availability (*A*) as:

$$A_{ij}^{(t)} = S\left(W_{i.}^{(t)}, C_{.j}^{(t)}\right).$$

Because regulation is the result of a gene being available to be regulated and a TF being responsible for the regulatory process, we average these values

$$\widetilde{W}_{ij}^{(t)} = \frac{1}{2}\left(A_{ij}^{(t)} + R_{ij}^{(t)}\right),$$

and use this to incrementally update the regulatory network *W* using a “learning rate” parameter, α (0<α<1) such that

$$W_{ij}^{(t+1)} = (1-\alpha)W_{ij}^{(t)} + \alpha\widetilde{W}_{ij}^{(t)}.$$

We take a similar approach to updating both the Co-regulatory (*C*) matrix for genes and the Cooperativity matrix for TFs (*P*), using our understanding of the regulatory process. For the Co-regulatory matrix, we reinforce our assumption that correlated expression should be tied to regulation by the same transcription factors. So, we define

$$\tilde{C}_{kj}^{(t)} = S\left(W_{.k}^{(t)}, W_{.j}^{(t)}\right),$$

and update *C* using

$$C_{kj}^{(t+1)} = (1-\alpha)C_{kj}^{(t)} + \alpha\tilde{C}_{kj}^{(t)}.$$

For the Cooperativity matrix (*P*), we reinforce our assumption that those TFs that form complexes should have genes with similar patterns of expression. Here we define

$$\tilde{P}_{im}^{(t)} = S\left(W_{i.}^{(t)}, W_{m.}^{(t)}\right),$$

and update *P* using

$$P_{im}^{(t+1)} = (1-\alpha)P_{im}^{(t)} + \alpha\tilde{P}_{im}^{(t)}.$$

Finally, at each iteration of network updates, we compute the Hamming distance (*H*) as a measure of convergence,

$$H^{(t)} = \left|\widetilde{W}^{(t)} - W^{(t-1)}\right| = \frac{1}{N}\sum_{i,j}\left|\widetilde{W}_{ij}^{(t)} - W_{ij}^{(t-1)}\right|,$$

where *N* is the number of possible edges in the network. This procedure repeats until the Hamming distance falls below 0.001.

#### 2. Similarity metrics

PANDA uses a similarity metric *S* to compute the pairwise agreement between three sources of biological evidence. By default, PANDA uses a modification of the Tanimoto similarity (Tfunction) as a similarity metric to account for continuous input:

$$t(x,y) = \frac{xy'}{\sqrt{xx' + yy' - |xy'|}},$$

with $x$ and $y$ representing vectors of size (1, n).

At each step in the iterative process, PANDA averages the availability matrix and the responsibility matrix. The responsibility matrix is obtained through computing $t(x,y)$ for each TF and gene pair with regards to the TF's relationships with all TFs in the PPI matrix (row in P represented by a vector $x$), and the TFs targeting the gene in the binding matrix (column in W represented by a vector $y$). The availability matrix is computed through computing $t(x,y)$ for each TF and gene pair with regards to the relationship between all the target genes of a TF (row in *W* represented by a vector *x*) and the target gene's coexpression with all genes in the coexpression matrix (column in *C* represented by a vector *y*).

This process is highly amenable to GPU acceleration, where each GPU core processes the computation of a single TF-gene similarity.

We extended PANDA to allow the use of a range of similarity metrics that were derived from the following distances. The Euclidean distance was computed as follows:

$$d_1(x,y) = \sqrt{(x-y)(x-y)'}$$

The squared Euclidean was computed as follows:

$$d_2(x,y) = (x-y)(x-y)'$$

The standardized Euclidean distance scales the Euclidean distance by the diagonal matrix $V_{(n,n)}$, that has the standard deviation of each vector as an entry.

$$d_3(x,y) = \sqrt{(x-y)V^{-1}(x-y)'}$$

The City block distance was computed as follows:

$$d_4(x,y) = \sum_{j=1}^{n} |x_j - y_j|$$

The Minkowski distance was computed as follows for p=3:

$$d_5(x,y) = \sqrt[p]{\sum_{j=1}^{n} |x_j - y_j|^{p}}$$

The Chebychev distance was computed as follows:

$$d_6(x,y) = max_j\{|x_j - y_j|\}$$

The Euclidean, Squared Euclidean, Standardized Euclidean, Squared Euclidean, City block, Minkowski, and Chebychev distances were converted to similarities as follows:

$$S = \frac{1}{1+d}$$

In addition, the following similarities were considered without further transformation. The cosine similarity was computed as follows:

$$s_1(x,y) = \frac{xy'}{(xx')(yy')}$$

The Pearson Correlation Coefficient was computed as follows:

$$s_2(x,y) = \frac{(x-y)(x-y)\prime}{\sqrt{(x-\overline{x})(x-\overline{x})\prime}\sqrt{(y-\overline{y})(y-\overline{y})\prime}} ,$$

with $\overline{x} = \frac{1}{n}\sum_j x_j$ and $\overline{y} = \frac{1}{n}\sum_j y_j$ .

#### II. Supplementary figures

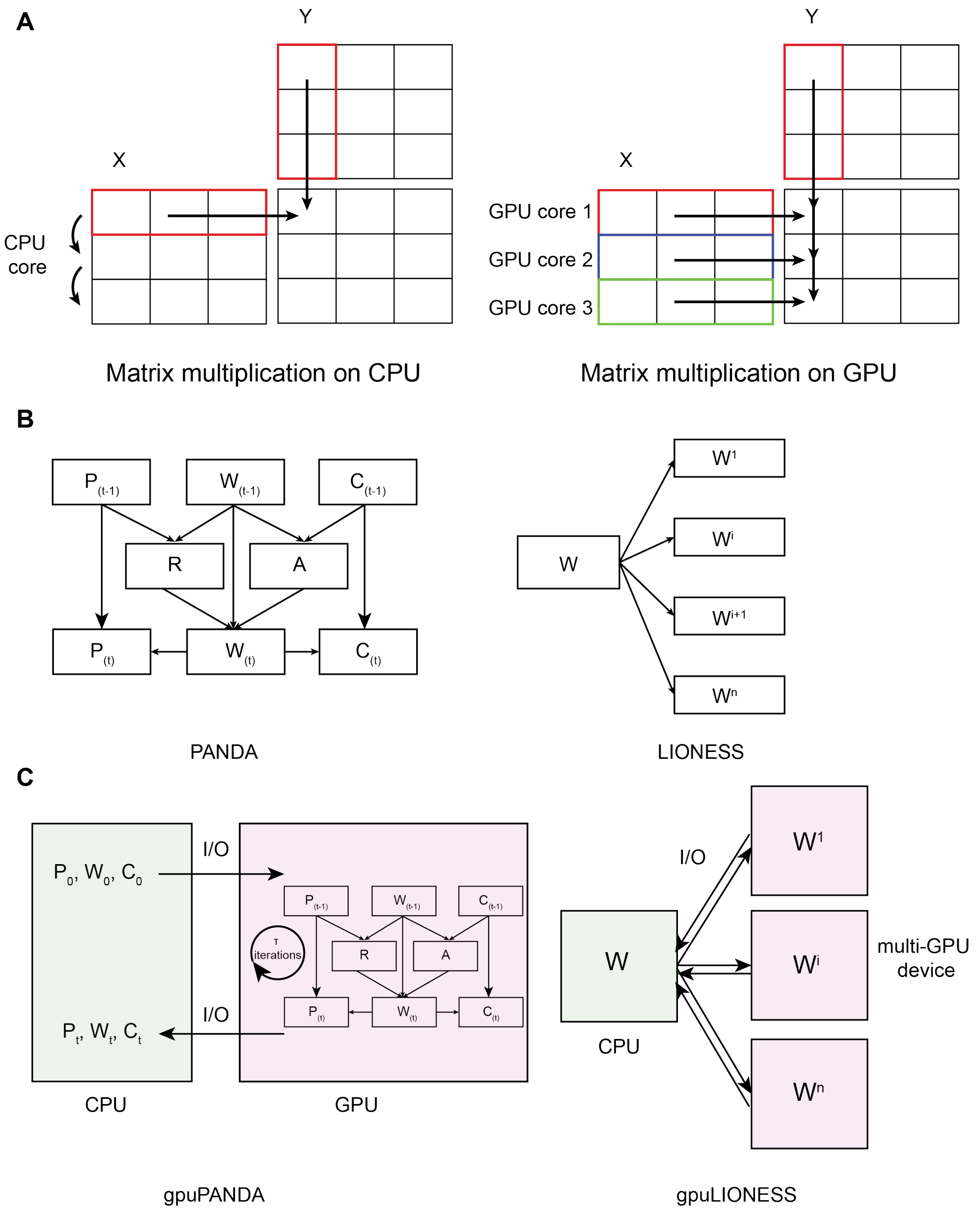


**Figure S1 – A schematic of GPU implementation in gpuZoo.** A - Matrix multiplication in GPU distributes elements-wise operations across many cores while CPU processes them sequentially. B - A representation of PANDA algorithm to generate a regulatory network *W* from three input networks $P_{(0)}$ for TF PPI, $C_{(0)}$ for gene coexpression, and $W_{(0)}$ the motif regulatory prior. LIONESS infers single-sample networks by linear interpolation from an input regulatory network. C - The gpuPANDA implementation starts by sending input variables to device memory, executes the iterative process on the device, and then retrieves the final results. gpuLIONESS parallelizes linear interpolation across several GPUs. A: Availability matrix, R: Responsibility matrix.

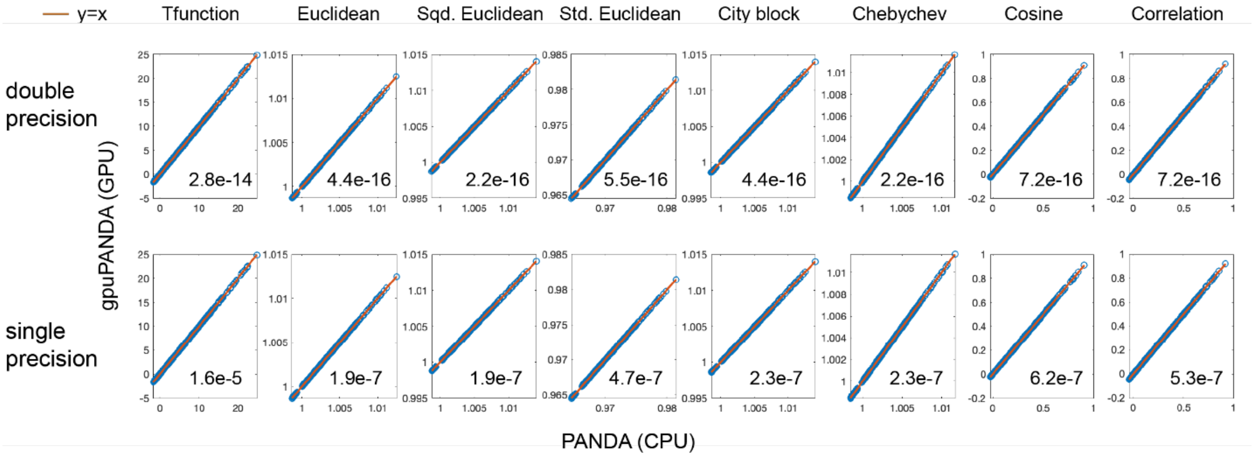


**Figure S2 - Comparison of small model edge weights between the GPU and CPU implementation of PANDA in single and double precision for eight similarity metrics.** The network edges (blue circles) are plotted against the unity line (orange line). The maximum absolute value of the difference between the networks is shown in the bottom right of each panel.

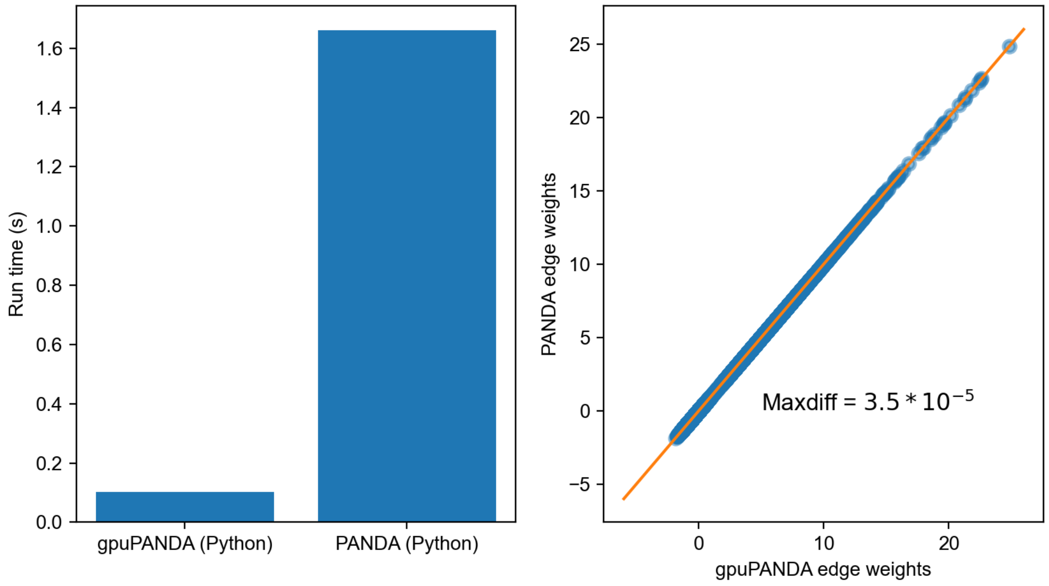


**Figure S3 – Performance of gpuPANDA in Python for the small model using GPU3, with alpha equal to 0.1 and using single precision computation.** Comparison of the runtime of gpuPANDA in Python using GPU3 and PANDA using GPU3 in seconds (left panel). Comparison of the edge weights between gpuPANDA in Python and PANDA using single precision plotted against the unity line (right panel). The largest absolute difference between the two networks is equal to $3.5 \times 10^{-5}$. Maxdiff: Maximum absolute difference.

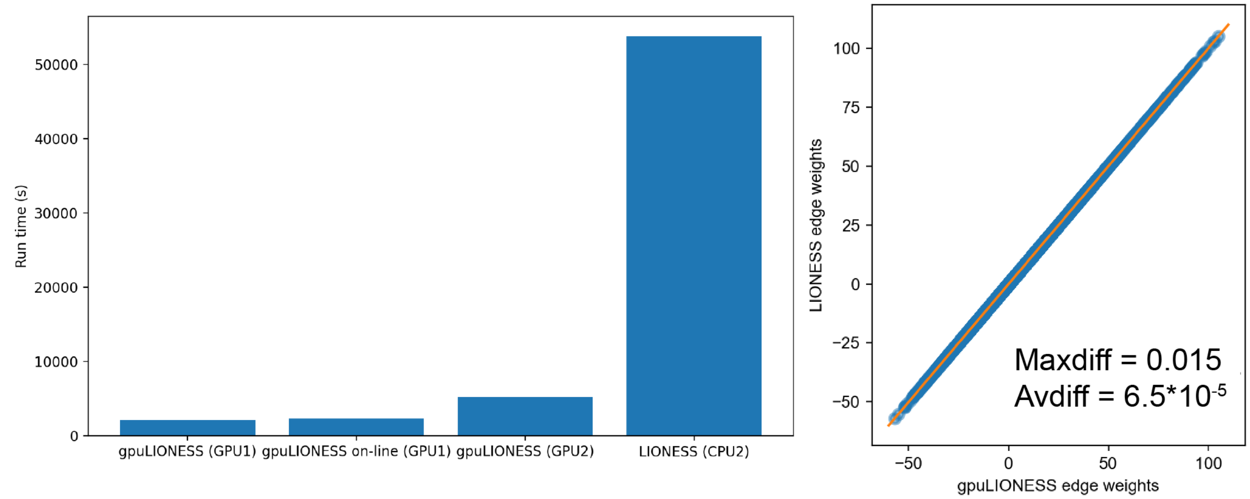


**Figure S4 - Performance of gpuLIONESS in GPU1 and GPU2 in comparison to CPU2 in coding gene networks in single precision.** Runtime of gpuLIONESS and LIONESS as implemented in MATLAB for the reconstruction of 127 sample-specific networks (left panel). Comparison of the edge weights between the LIONESS and gpuLIONESS networks for the first sample with the largest absolute difference equal to 0.015 and the average absolute difference equal to $6.5x10^{-3}$. Maxdiff: Maximum absolute difference, Avdiff: Average absolute difference.

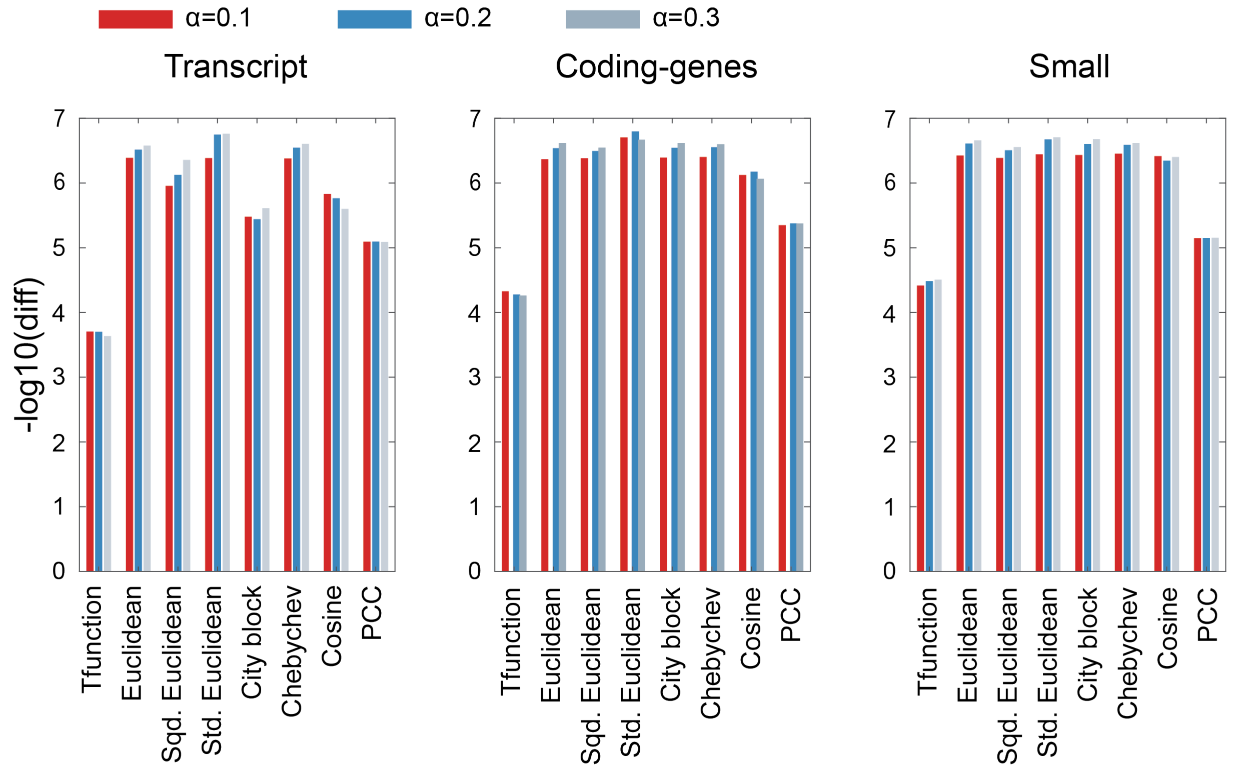


**Figure S5 - Comparison of single precision and double precision networks.** We compared the difference between transcript, protein-coding genes, and small-sized networks in single and double precision computed in CPU across three learning rate values and eight similarity metrics.

The y axis represents the -log10 of the largest difference between the single and double precision networks.

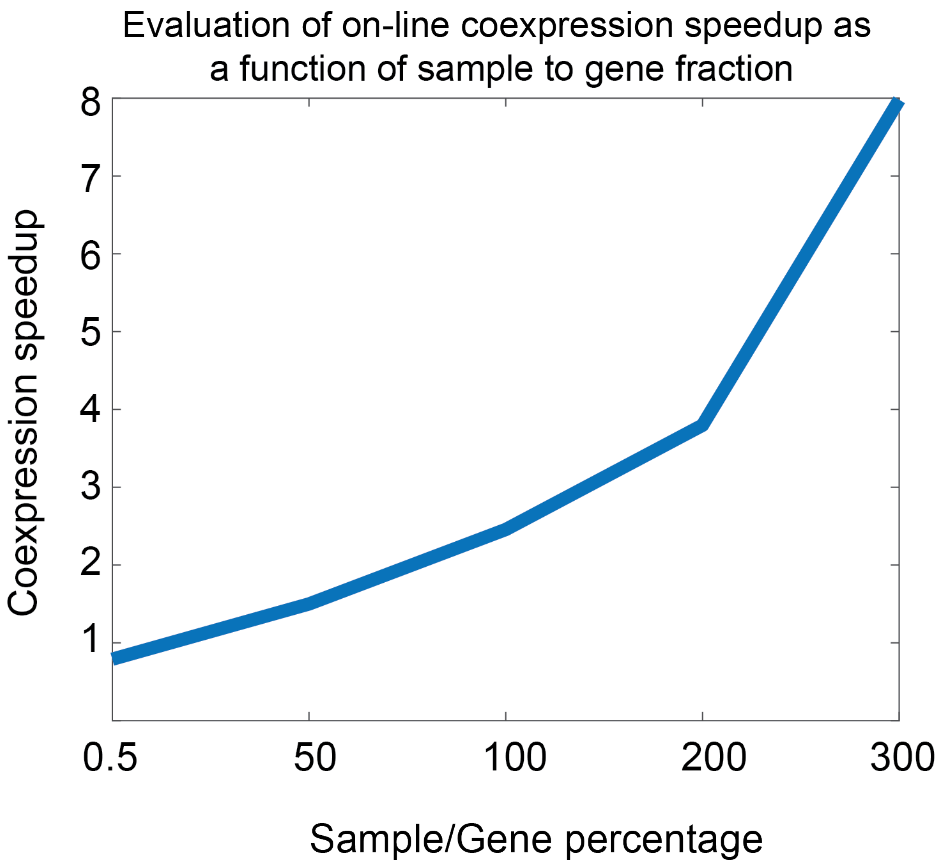


**Figure S6 – Evaluation of on-line gene coexpression speedup as a function of the sample to genes percentage.** Speedup was computed as fraction of CPU time over GPU time.

#### III. Supplementary table legends

**Table S1 – Runtime fold change of MATLAB gpuPANDA on CPU1/GPU1, CPU2/GPU1, CPU2/GPU2, CPU1/GPU3, and CPU2/GPU3, and CPU1/GPU2.** Empty values mean that either gpuPANDA could not load the data on device memory or the simulation did not finish in reasonable time. The highlighted entries indicate the metric that has the largest speedup in GPU vs CPU for a given value of alpha.

**Table S2 – Runtime of MATLAB gpuPANDA on GPU1, GPU2, GPU3, CPU1, and CPU2.** Empty values mean that either gpuPANDA could not load the data on device memory or the simulation did not finish in reasonable time.

**Table S3 – Computing cost fold change of MATLAB gpuPANDA on CPU1/GPU1, CPU2/GPU1, CPU2/GPU2, CPU1/GPU3, and CPU2/GPU3, and CPU1/GPU2.** Empty values mean that either gpuPANDA could not load the data on device memory or the simulation did not finish in reasonable time. The highlighted entries indicate the metric that has the largest cost decrease in GPU vs CPU in each alpha.

**Table S4 – Computing cost of MATLAB gpuPANDA on GPU1, GPU2, GPU3, CPU1, and CPU2.** Empty values mean that either gpuPANDA could not load the data on device memory or the simulation did not finish in reasonable time.
